# The antiparasitic drug atovaquone inhibits arbovirus replication through the depletion of intracellular nucleotides

**DOI:** 10.1101/507152

**Authors:** Angelica Cifuentes Kottkamp, Elfie De Jesus, Rebecca Grande, Julia A. Brown, Adam R. Jacobs, Jean K. Lim, Kenneth A. Stapleford

## Abstract

Arthropod-borne viruses represent a significant public health threat worldwide yet there are few antiviral therapies or prophylaxis targeting these pathogens. In particular, the development of novel antivirals for high-risk populations such as pregnant women is essential to prevent devastating disease such as that which was experienced with the recent outbreak of Zika virus (ZIKV) in the Americas. One potential avenue to identify new and pregnancy friendly antiviral compounds is to repurpose well-known and widely used FDA approved drugs. In this study, we addressed the antiviral role of atovaquone, a FDA Pregnancy Category C drug and pyrimidine biosynthesis inhibitor used for the prevention and treatment of parasitic infections. We found that atovaquone was able to inhibit ZIKV and chikungunya virus virion production in human cells and that this antiviral effect occurred early during infection at the initial steps of viral RNA replication. Moreover, we were able to complement viral replication and virion production with the addition of exogenous pyrimidine nucleosides indicating that atovaquone is functioning through the inhibition of the pyrimidine biosynthesis pathway to inhibit viral replication. Finally, using an *ex vivo* human placental tissue model, we found that atovaquone could limit ZIKV infection in a dose-dependent manner providing evidence that atovaquone may function as an antiviral in humans. Taken together, these studies suggest that atovaquone could be a broad-spectrum antiviral drug and a potential attractive candidate for the prophylaxis or treatment of arbovirus infection in vulnerable populations, such as pregnant women.

**Author Summary:** The ability to protect vulnerable populations such as pregnant women and children from Zika virus and other arbovirus infections is essential to preventing the devastating complications induced by these viruses. One class of antiviral therapies may lie in known pregnancy-friendly drugs that have the potential to mitigate arbovirus infections and disease yet this has not been explored in detail. In this study, we show that the common antiparasitic drug, atovaquone, inhibits arbovirus replication through intracellular nucleotide depletion and can impair ZIKV infection in an *ex vivo* human placental explant model. Our study provides a novel function for atovaquone and highlights that the rediscovery of pregnancy-acceptable drugs with potential antiviral effects can be the key to better addressing the immediate need for treating viral infections and preventing potential birth complications and future disease.

## Introduction

Recent outbreaks of significant human vector-borne pathogens have left us with the uncertainty of potential future devastating epidemics [1], [2]. In particular, Zika virus (ZIKV), a flavivirus and close relative of dengue virus (DENV), has led to an overwhelming spectrum of diseases, including Guillain-Barré syndrome, microcephaly, ocular and testicular damage, and even meningitis, encephalitis, thrombocytopenia and multiorgan failure [3], [4], [5], [6], [7], [8]. This, coupled with the widespread and invasive *Aedes* species of mosquito makes it easy to envision another epidemic when environmental, ecological, and human factors meet [9]. Unfortunately, there are no antiviral treatments or prophylaxes targeting these viruses, and thus efforts to mitigate and ultimately prevent the impact of the disease are urgent and need to be addressed.

Pregnant women carry a particularly high risk for ZIKV and other arbovirus-related complications [10], [11], [12]. Importantly, the capacity of the virus to infect trophoblasts, Hofbauer macrophages and endothelial cells [1], [13], thus allowing it to infect the fetus at any stage of growth, challenges the protective function of the placenta in the materno-fetal interface [14],[15]. Despite the significant morbidity observed in newborns [16], there are no antivirals available to treat this population in part due to safety concerns during pregnancy, lack of biosafety studies and nonexistent clinical trials. With this in mind, and given the urgency of this need, we propose to repurpose existing drugs with an acceptable profile in pregnancy.

Nucleotide biosynthesis inhibitors such as ribavirin, brequinar, and mycophenolic acid (MPA) have been shown extensively to inhibit a wide array of viral infections both *in vitro* and *in vivo* [17], [18], [19], [20], [21] [22], [23]. In addition, a number of small compounds that possess antiviral function through the depletion of intracellular nucleotide pools have been identified, suggesting that this cellular pathway may be a prime target for antiviral development [24], [25], [26], [27], [28]. Unfortunately, many of these compounds have numerous side effects and are not approved for use in high risk populations such as pregnant women or children, thus the development of safe and pregnancy-acceptable nucleotide biosynthesis inhibitors would be ideal candidates as antivirals.

In these studies, we address the antiviral role of atovaquone, a FDA Pregnancy Category C and well-known antimalarial and antiparasitic drug that has been used repeatedly in the clinical setting for nearly two decades [29] [30], [31], [32]. Atovaquone is a ubiquinone (Coenzyme Q) analogue that functions through the inhibition of the mitochondrial cytochrome complex III [33, 34]. However, it has also been shown to inhibit dihydroorotate dehydrogenase (DHODH), an enzyme required for *de novo* pyrimidine synthesis, leading to specific depletion of intracellular nucleotide pools [33], [35], [36], [37]. Given these capacities, we hypothesized that atovaquone may function similarly to other known nucleotide biosynthesis inhibitors and may inhibit RNA virus replication.

Here, we show that atovaquone is able to inhibit ZIKV and chikungunya virus (CHIKV) replication and virion production in human cells, similar to what has been shown for other pyrimidine biosynthesis inhibitors. Moreover, we found this effect to occur early in infection, during the initial steps of viral RNA synthesis and that viral inhibition can be rescued with the addition of exogenous pyrimidines, indicating this drug is functioning through the blocking of DHODH and depletion of intracellular nucleotides. Finally, we show that atovaquone can inhibit ZIKV infection in an *ex vivo* human placental tissue model. Taken together, these studies identify atovaquone as a potential pregnancy acceptable antiviral compound. More importantly, they highlight the potential to repurpose available drugs in the hopes to one day translate these findings to novel and safe approaches, preventing arbovirus-related outcomes of vulnerable populations.

## Results

### Atovaquone inhibits arbovirus replication *in vitro*

Nucleotide biosynthesis inhibitors have been shown to have antiviral activity towards a wide range of RNA viruses both *in vitro* and *in vivo* [18], [21], [26], [38], [39], [40], suggesting that manipulating this pathway is a potential avenue for antiviral development. However, many of these compounds are not approved for use in high-risk populations such as pregnant women. Atovaquone is a well-tolerated antiparasitic [31] drug that has been used extensively for the treatment and prevention of *pneumocystis jirovecii* pneumonia (PCP), toxoplasmosis, babesiosis and malaria [29], [30], [32], [34], yet the antiviral role of atovaquone has not been addressed. To examine the antiviral activity of atovaquone, we pretreated Vero cells with increasing concentrations of atovaquone as well as ribavirin, MPA, and brequinar, known nucleotide biosynthesis inhibitors shown to have antiviral function (**Figure 1**). The cells were subsequently infected with either the Ugandan or Brazilian strains of Zika virus (ZIKV) and viral inhibition was assessed 72 hours post infection by immunostaining for the ZIKV envelope (E) protein. We found that all known nucleotide biosynthesis inhibitors were able to inhibit ZIKV replication although in a strain specific manner (**Figure 1A-C**). Importantly, we found that atovaquone exhibited similar antiviral activity over the concentrations tested and that this inhibition was again strain specific (**Figure 1D-F**). Taken together, these studies show that atovaquone inhibits ZIKV infection and spread, potentially through the depletion of intracellular nucleotide pools.

**Figure 1:**
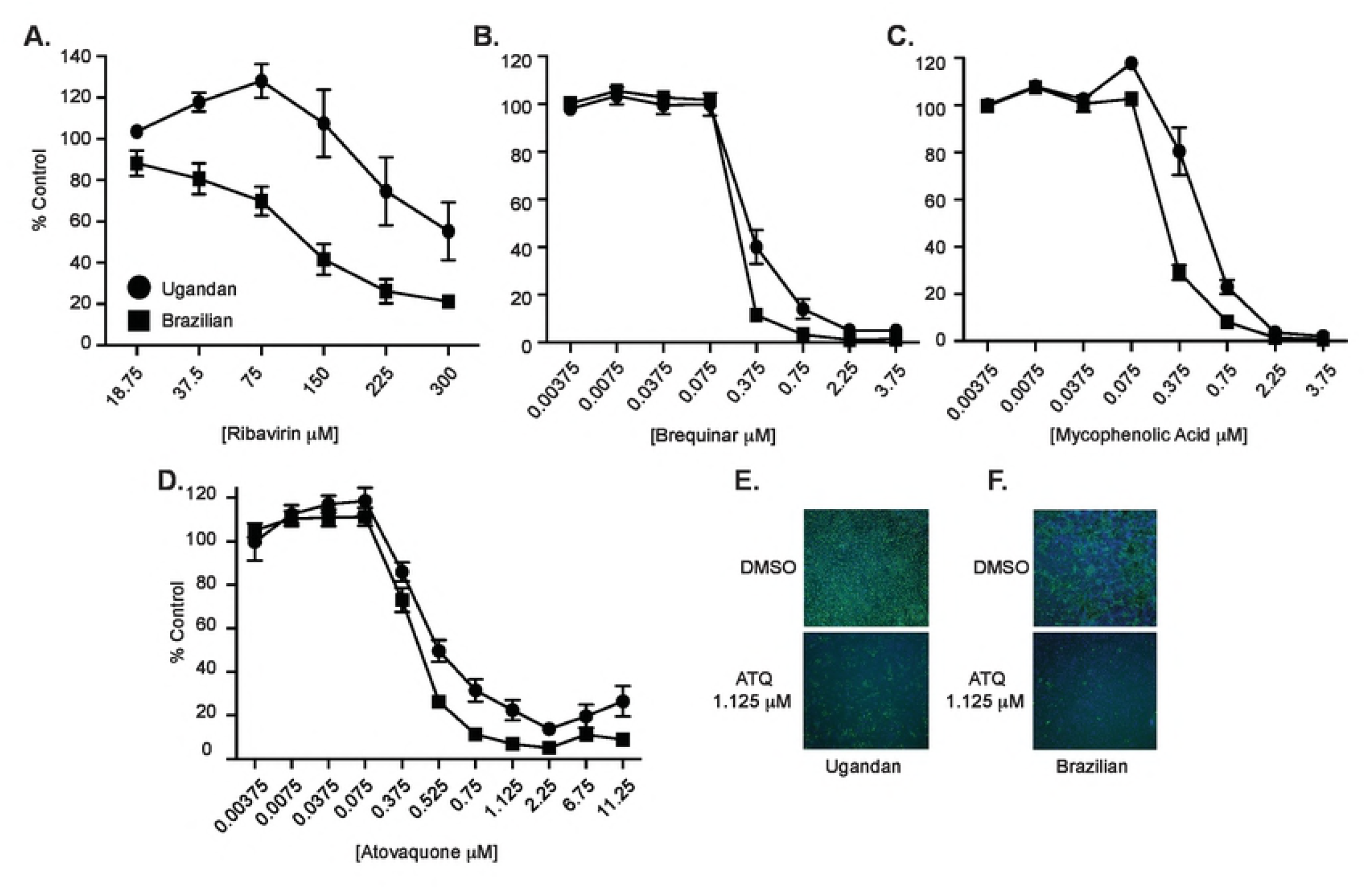
Nucleotide biosynthesis inhibitors impair ZIKV replication. Vero cells were pretreated with (**A**) ribavirin, (**B**) brequinar, (**C**) mycophenolic acid, (**D**) atovaquone or carrier controls for two hours and subsequently infected with either the Ugandan (MR766) or Brazilian (Paraiba_01/2015) strain of ZIKV at a MOI of 0.1. Cells were fixed 72 hours post infection, stained with anti-flavivirus E antibody, and infected cells were quantified on a CellInsight CX7 high-content microscope. Data are represented as percent ZIKV positive cells compared to the carrier control. (**E and F**) Representative images of Ugandan (**E**) and Brazilian (**F**) atovaquone inhibition at 1.125 μM.

### Atovaquone impairs ZIKV virion production in human cells

Given the inhibition of viral replication in Vero cells, we next addressed whether atovaquone was able to reduce the production of infectious ZIKV particles in mammalian cell types including human cells. For these and all subsequent studies, we chose to use the Ugandan strain of ZIKV due to its robust replication *in vitro* and its relative resistance to atovaquone compared to the Brazilian strain (**Figure 1**). We infected Vero, human 293T and human placental JEG3 cells with the Ugandan strain of ZIKV in the presence of atovaquone or a DMSO control and quantified infectious virion production by plaque assay. We found that atovaquone significantly impaired virion production in all cell types tested although the peak of inhibition varied between cell type (**Figure 2A-C**).

One potential explanation for these results could be that atovaquone is toxic, and this leads to reduced virus production. To address this, we first measured cell viability by a MTT cell proliferation assay and found that atovaquone indeed had a dose-dependent reduction in cell proliferation compared to the DMSO control (**Figure S1A**). However, atovaquone is a mitochondrial cytochrome complex III inhibitor [33, 36, 44] and thus we hypothesized that the MTT assay results we observed may be due to mitochondrial inhibition and are not directly indicative of dying cells. To confirm this, we measured cell viability with Sytox Green, a dye that binds nucleic acids when both the plasma and the nuclear membrane are permeabilized and thus represents dying cells (**Figure S1B**). Using this assay, we found that high concentrations of atovaquone, particularly in 293T cells, did lead to more cell death than the DMSO control; however, lower concentration, that did show effects by the MTT assay, had minimal effects on cell viability. Moreover, these results were confirmed in our data as although we observed a reduction in cell growth by MTT assay we found that in Vero and 293T cells, virus production increased at higher concentrations of atovaquone suggesting that the cells are still competent for virus production under these conditions. Taken together, these results show that atovaquone is able to inhibit ZIKV virion production in mammalian and human cell types.

To expand on these findings, we addressed if atovaquone could inhibit chikungunya virus (CHIKV), an arbovirus transmitted by *Aedes* mosquitoes and capable of causing severe human disease and significant outbreaks [45], [46], [47]. Importantly, CHIKV has been shown to be inhibited by nucleotide biosynthesis inhibitors [22], [48], [49], and thus we hypothesized it would be inhibited by atovaquone. We infected Vero cells with a CHIKV expressing a ZsGreen reporter in the presence of increasing concentrations of atovaquone and quantified the number of ZsGreen positive cells after treatment by microscopy. Similar to ZIKV, CHIKV replication was inhibited by atovaquone in a dose-dependent manner (**Figure 3A**). This inhibition in replication was further confirmed in Vero and 293T cells where we observed reductions in infectious CHIKV virions after treatment with atovaquone (**Figure 3B**). These results suggest that atovaquone can inhibit multiple arboviruses and has the potential to be used as a well-tolerated antiviral therapy.

### Atovaquone inhibits ZIKV at early stages of infection

To explore which stage of viral life cycle atovaquone targets, we first addressed viral entry by treating cells with atovaquone or DMSO during virus entry, washing the cells extensively, and adding back complete media (**Figure 4A**, at entry). As a control, we added atovaquone during entry and added back media containing atovaquone after infection (**Figure 4A**, post entry). We found that when cells are treated during viral entry there is no change in production of viral particles compared to DMSO yet when added after entry atovaquone was able to inhibit ZIKV replication suggesting that atovaquone functions post entry. To investigate this further, we performed time-of-addition experiments to test which part of the intracellular viral life cycle is targeted by atovaquone. We infected 293T cells with ZIKV, added atovaquone or a DMSO control at different time-points during infection (**Figure 4B**) and quantified viral titers by plaque assay at 36 hours post-infection. We found that ZIKV virion production is most inhibited at early stages of infection (up to 4 hours post infection) and that this effect diminishes as the infection progresses, suggesting that atovaquone acts early during infection, potentially inhibiting RNA replication. Finally, to investigate the impact of atovaquone on ZIKV RNA replication, we analyzed ZIKV intracellular RNA at multiple time points post infection (**Figure 4C**). We found that where viral RNA levels were equal after infection (time 0), again confirming there is no effect of atovaquone on viral entry, there was a significant difference in viral RNA at 24 and 48 hours post infection indicating that atovaquone is acting to inhibit ZIKV RNA replication early during infection.

### ZIKV RNA replication and virion production is rescued by the addition of exogenous nucleosides

Given the role of atovaquone in the inhibition of DHODH and similar viral inhibition curves to brequinar, another inhibitor of DHODH and pyrimidine biosynthesis (**Figure 1B**), we hypothesized that atovaquone may function through a similar pathway. To address this, we performed a rescue experiment where we infected cells with ZIKV in the presence of atovaquone or DMSO followed by media with atovaquone supplemented with 100 μM uridine, cytidine, adenosine, or guanosine. We found that in all cells types, ZIKV infectious particle production was rescued only when uridine was added to the media (**Figure 5A-C**). Given these results and the dual function of atovaquone in blocking both mitochondrial function and DHODH, it is possible that the addition of exogenous nucleosides could simply have rescued the MTT phenotype (mitochondrial function) we see for atovaquone and thus ZIKV replication. However, we found that when cells were incubated in the presence of atovaquone and nucleoside there was no change in MTT cell proliferation (**Figure S2**), indicating that the ZIKV inhibition and rescue we observe is not through mitochondrial inhibition but rather through the inhibition of DHODH. Furthermore, we addressed the ability of uridine to complement ZIKV RNA synthesis and found that indeed the addition of exogenous uridine rescued this phenotype to similar levels of the DMSO control (**Figure 5D**). Taken together, these results suggest that atovaquone is functioning through the depletion of intracellular nucleotide pools, and the addition of exogenous uridine can rescue ZIKV replication via the pyrimidine salvage pathway, bypassing the inhibition of atovaquone at critical steps in the *de novo* pyrimidine synthesis inhibition.

### Atovaquone inhibits ZIKV infection in an *ex vivo* human placental tissue model

We found that atovaquone significantly inhibited ZIKV in human placental JEG3 cells *in vitro* and thus were interested in determining the extent to which this compound could inhibit ZIKV infection in an *ex vivo* human placental tissue model. To investigate this, we infected human placental chorionic villus explants with the Ugandan strain of ZIKV in the presence of increasing doses of atovaquone. Similar to data in cell lines, we found that ZIKV infection and virion production were inhibited in a dose-dependent manner in the human placental tissue (**Figure 6A**). These results were confirmed by fluorescence *in situ* hybridization probing for ZIKV RNA in ZIKV infected tissue where atovaquone treatment reduced ZIKV spread (**Figure 6B**), showing a dose-dependent decrease in ZIKV-infected cells in the presence of increasing amounts of atovaquone. These data highlight that atovaquone may provide protection in the human placenta-fetal interface during ZIKV infection.

## Discussion

It remains unknown when the next outbreak of ZIKV will occur, yet we know through past devastating epidemics, in which thousands of women and children were affected by this virus that we still have an urgent need for effective therapies against ZIKV infection [11], [50], [51]. Despite current potential protection from herd and self-immunity, environmental factors, and host-vector-virus interactions that keep ZIKV in the low incidence figures [1], [52], [53], [54], [55], preventing ZIKV and other arbovirus infections should be a priority. In recent years, many compounds have been proposed as potential anti-ZIKV agents following *in vitro* results [24], [25], [56], [57], [58], [59], [60], [61], [62], [63], [64], [65]. Some of these drugs have an extensive background in the medical field and offer attractive options, either alone or in combination, for treatment and perhaps prophylaxis of ZIKV infections; however, most of them remain inadequate to be used during pregnancy. Only chloroquine has been demonstrated in pregnancy animal models to be effective against ZIKV [56] [57] and none of them have been tested in humans in the context of ZIKV infection. Here, we propose a pregnancy-acceptable drug candidate for the treatment of ZIKV and other potential viral infections, highlighting the repurposing of FDA approved drugs as a possible avenue for antiviral development.

Atovaquone, a ubiquinone analogue approved in humans since 1999 for the treatment of *Pneumocystis jiroveci* pneumonia (PCP) [32], [34] and prevention of malaria [29], has no antiviral activity described in the literature to date. However, given that atovaquone functions through the inhibition of pyrimidine biosynthesis, a pathway essential to viruses and the target of numerous antiviral compounds, we hypothesized that atovaquone may be antiviral as well. In this work, we addressed the antiviral role of atovaquone on ZIKV infections *in vitro* and in an *ex vivo* human placental tissue model, as well as explored the antiviral effect on CHIKV *in vitro.* We first screened atovaquone and the known nucleotide biosynthesis inhibitors and antiviral compounds, ribavirin, MPA, and brequinar, for their ability to inhibit two genetically distant strains of ZIKV. We found that ribavirin, MPA, and brequinar were able to inhibit ZIKV replication as has been shown previously and that atovaquone also led to inhibition of ZIKV replication behaving similarly to the pyrimidine biosynthesis inhibitor brequinar. Interestingly, we found that there was a strain-specific impact of the nucleotide biosynthesis inhibitors suggesting that genetic differences between the Ugandan and Brazilian strains may be responsible for the sensitivity to nucleotide depletion. Nonetheless, we concluded that atovaquone inhibits ZIKV replication similarly to other nucleotide biosynthesis inhibitors in Vero cells.

One potential caveat of these experiments could be that whereas we do detect reductions in ZIKV infected cells by immunofluorescence, this could have little impact on infectious virion production. To address this, we quantified infectious virus in the supernatant in the presence of atovaquone treatment compared to a DMSO control. As we saw with immunofluorescence, atovaquone was able to significantly reduce the amount of infectious virus in multiple mammalian cell types, including human placental cells over a subset of drug concentrations. Interestingly we found that in Vero and 293T cells, higher concentrations of drug had the least impact on virion production. One explanation for this could be that the virus has evolved to be resistant to atovaquone. However, we find this unlikely given the short time of infection and that drug resistance typically takes multiple passages. An additional explanation could be that at high concentrations, atovaquone is interfering with other cellular pathways that impede its antiviral effects yet still results in the inhibition of mitochondrial function *in vitro.* This may be of particular importance for the use of atovaquone as an antiviral in humans. In the *in vitro* human cell culture system, the CC_50_ of atovaquone was roughly 10 μM which contrasts with historical trials of atovaquone taken orally at a doses of 750 mg every 6 hours for the treatment of toxoplasmosis, reporting steady serum concentrations in humans of roughly 50 μM without associated toxicity [36] [66] [67]. One possible explanation for this difference is that *in vitro* studies do not entirely represent all the biological interactions that take place in the human body. In addition, it is possible that at this high concentration, atovaquone is not antiviral and thus would need to be optimized at lower concentrations for its antiviral function in humans.

To address the stage in the viral life cycle where atovaquone acts, we performed viral entry assays by infecting cells in the presence or absence of atovaquone and then adding media with or without the compound after infection. Here, we found that adding atovaquone at the time of infection had no effect on virion production whereas adding atovaquone after infection was able to reduce ZIKV replication, indicating that atovaquone is acting post entry. We then performed time-of-addition assays where we added atovaquone one hour before, during, and at multiple time points post infection. We found that only early during infection, within the first two to four hours post infection, was atovaquone able to inhibit viral replication. These data are similar to what has been seen for the antiviral effects of brequinar on DENV [20], [41], suggesting atovaquone may function by a similar mechanism. Given these results, we hypothesized that atovaquone is functioning during the initial steps of ZIKV RNA replication and blocking virion production through the inhibition of RNA replication. When we quantified ZIKV RNA levels over time we found that indeed atovaquone treatment significantly reduced viral RNA synthesis.

Atovaquone is thought to function primarily through the inhibition of mitochondrial cytochrome complex III and thus inhibition of mitochondrial function in the parasite [34], [36], [37]. Using an MTT assay of cell proliferation measured through mitochondrial function we also saw that atovaquone was able to reduce mitochondrial function in all cell lines we analyzed, yet these cells were shown to be viable by Sytox Green staining. Atovaquone also functions through inhibiting dihydroorotate dehydrogenase (DHODH) [33], an enzyme involved in pyrimidine biosynthesis and in particular the synthesis of uridine monophosphate (UMP). Given the striking similarities to brequinar, another pyrimidine biosynthesis inhibitor, and that atovaquone specifically inhibited ZIKV RNA synthesis we hypothesized that this inhibition was through intracellular nucleotide depletion. To address this, we added exogenous nucleosides in the presence of atovaquone and indeed found that the addition of uridine was able to rescue ZIKV infection in all cell types, although to various extents. Moreover, we found that the addition of uridine was able to specifically rescue ZIKV RNA synthesis indicating that the antiviral effects of the drugs are through the depletion of intracellular nucleotides. We found it interesting that human 293T and JEG3 cells were unable to be completely rescued with the addition of uridine. However, it has been shown that nucleotide depletion will induce an antiviral innate immune response [26], [27], [40], and we hypothesize that a similar mechanism may be induced in these cells allowing them to retain antiviral function in the presence of exogenous uridine. One striking finding was that atovaquone in the presence of cytidine had an additive antiviral effect in several cell lines suggesting again that multiple cellular pathways may be at play. Finally, we show that atovaquone also works to inhibit ZIKV infection in human first and second trimester placental explants. Many studies have shown that placental cytotrophoblasts, trophoblasts, syncytotrophoblasts, Hofbauer cells, and fibroblasts are susceptible to ZIKV [13], [15], [68], [69], [70], [71], [72]; thus recreating the dynamics of this infection and host-virus interaction at the placental level makes this study relevant to the most vulnerable target of ZIKV, pregnant women.

Taken together, we found that atovaquone, a pregnancy-acceptable and common antiparasitic drug, has antiviral activity against ZIKV and CHIKV which can be translated in the clinical setting into an attractive candidate for the treatment and prevention of arbovirus infections in vulnerable populations as well as in individuals who live in or travel to endemic areas. Furthermore, patients with AIDS, chronic steroid users, and post-transplant patients that take atovaquone daily for the prevention of PCP, remain at high risk of acquiring multiple viral infections due to their impaired immune system. Therefore, it could be valuable to estimate the effect of atovaquone in viruses relevant to these patients such as human cytomegalovirus [73], herpes simplex virus [74], JC virus [75], and respiratory syncytial virus [76]. Atovaquone (in combination with proguanil hydrochloride) is already commercially available and broadly prescribed for malaria prophylaxis, yet so far there have not been proposed any clinical trials that address the relationship of atovaquone and ZIKV or CHIKV infections. This raises several questions: *i)* Are individuals who are/were taking atovaquone-proguanil (Malarone®) protected from viral threats as well? and *ii)* Does the broad administration of drugs which may unknowingly possess antiviral functions impact the evolution of viral infections? Future studies addressing these questions will be essential to understanding the antiviral function of atovaquone on viral evolution and disease. Nonetheless, these results contribute to the urgent need of finding effective ZIKV treatments especially for pregnant women, as these treatments should be readily accessible in order to ameliorate the teratogenic consequences of ZIKV across all trimesters of pregnancy. The studies completed here identified a potential candidate for these at-risk populations, yet more work is needed to define the complete antiviral role of atovaquone *in vivo.* Moreover, these studies have highlighted that repurposing drugs may provide fast avenues to the development of novel antiviral therapies and that we can potentially exploit FDA approved, pregnancy-friendly drugs to fight emerging viral threats.

## Materials and Methods

### Cells and viruses

Vero cells (CCL-81, ATCC) were cultured in Dulbecco Modified Eagle Medium (DMEM) (Corning) supplemented with 10% new born calf serum (NBCS, Gibco), 100 μg/mL penicillin-streptomycin (P/S) (Corning) at 37°C with 5% CO_2_. BHK-21 (CCL-10, ATCC), 293T (CRL-3216, ATCC), and JEG3 (provided by Dr. Carolyn Coyne, University of Pittsburgh) were cultured in DMEM supplemented with 10% fetal bovine serum (FBS, Atlanta Biologicals), 1% nonessential amino acids (NEAA, Corning), and 1% P/S at 37°C with 5% CO_2_. All cell lines were confirmed to be mycoplasma free.

The Ugandan (MR766) [77] and Brazilian (Paraiba_01/2015) [78] strains of ZIKV were generated from [79] infectious clones provided by Dr. Matthew Evans (Icahn School of Medicine at Mount Sinai) and Dr. Alexander Pletnev (National Institutes of Health), respectively. To generate initial viral stocks, each plasmid was transfected via lipofectamine 2000 reagent (Invitrogen) into 293T cells and virus containing supernatant was harvested 48 hours post transfection. A working viral stock was then generated by passaging the initial viral stock over Vero cells. Viral titers were quantified by plaque assay. In brief, 10-fold serial dilutions of each virus in DMEM were added to a confluent monolayer of Vero cells for 1 hour at 37°C. Following incubation, cells were overlaid with 0.8% agarose in DMEM and 2% NBCS and incubated at 37°C for five days. The cells were fixed with 4% formalin, agarose plugs were removed, and plaques were visualized by crystal violet.

Wild type CHIKV was generated from the La Reunion 06-049 infectious clone as previously described [79] [48]. A CHIKV La Reunion infectious clone expressing ZsGreen was constructed by standard molecular biology techniques. First, an AvrII restriction enzyme site was inserted 5’ of the subgenomic promoter by site-directed mutagenesis using the primers (Forward 5’ CACTAATCAGCTACACCTAGGATGGAGTTCATCCC 3’ and Reverse 5’ GGGATGAACTCCATCCTAGGTGTAGCTGATTAGTG 3’). The CHIKV subgenomic promoter was then amplified by PCR (Forward 5’ CCTAGGCCATGGCCACCTTTGCAAG 3’ and Reverse 5’ ACTAGTTGTAGCTGATTAGTGTTTAG 3’) and subcloned into the AvrII site to generate a CHIKV infectious clone containing two subgenomic promoters. Finally, the ZsGreen cassette was amplified by PCR (Forward 5’ GTGTACCTAGGATGGCCCAGTCCAAGCAC 3’ and Reverse 5’ GCTATCCTAGGTTAACTAGTGGGCAAGGC 3’) from a CHIKV infectious clone obtained from Dr. Andres Merits (University of Tartu) and subcloned into the AvrII restriction enzyme site. The complete cassette and subgenomic regions were sequenced to ensure there were no second-site mutations. To generate infectious virus, each plasmid was linearized overnight with NotI, phenol-chloroform extracted, ethanol precipitated, and used for *in vitro* transcription using the SP6 mMessage mMachine kit (Ambion). *In vitro* transcribed RNAs were phenol-chloroform extracted, ethanol precipitated, aliquoted at 1 μg/μl, and stored at −80°C. 10 μg of each RNA was electroporated into BHK-21 cells [23] and virus was harvested 48 hours post electroporation. Working virus stocks were generated by passaging virus over BHK-21 cells and viral titers were quantified by plaque assay as described above.

### Ethics Statement

For these experimental studies, first and second trimester human placental tissue was obtained within two hours of surgery from donors undergoing elective termination under an IRB protocol approved by the Institutional Review Board for the Ichan School of Medicine at Mt. Sinai (HS: 12-00145). All subjects provided informed written surgical consent for the use of de-identified waste materials for educational research. Tissue specimens are considered to be non-human subjects since they are de-identified. Following the surgery, tissue specimens are delivered to Mount Sinai’s Institutional Biorepository and Molecular Pathology Shared Resource Facility (SRF) in the Department of Pathology. The biorepository operates under a Mount Sinai Institutional Review Board (IRB) approved protocol and follows guidelines set by HIPAA. All samples are linked, with appropriate IRB approval and consent, to clinical and pathological data, and are open to all investigators of the institution, as well as to specific third-party collaborative efforts with investigators from other institutions.

### *Ex vivo* infection of human placental tissue

First and second trimester human placental tissue was obtained as described above. Chorionic villi adjacent to the fetal chorionic plate were placed into prewarmed DMEM containing 25% F-12 media, 10% FBS, 5 mM HEPES, 2 mM Glutamine, 100 IU/ml Penicillin, 100 μg/ml Streptomycin, 2.5 μg/ml Fungizone, and 300 ng/ml Timentin as previously described [80]. After removal of the amnion and decidua, the chorionic villi were cut into 0.2 cm^3^ blocks; nine blocks were plated per well of a six-well plate onto collagen gelfoams (Cardinal Health) in 3 ml media, and 3 wells were used per condition (27 tissue blocks). Following an overnight incubation at 37°C, tissue blocks were individually infected with 1×10^5^ PFU ZIKV^MR766^ in a volume of 5 μl that was pre-incubated with 15 μM, 5 μM, or 1.6 μM atovaquone for 1 hour. Atovaquone was maintained in the culture media at the same concentrations throughout 6 days of culture. Supernatants were collected and media changed every other day.

### ZIKV plaque forming unit assay

ZIKV plaque forming units (PFU) were quantified on Vero cell monolayers whereby 250 μl of tissue culture supernatant was adsorbed for 2 hours at 37°C in 12 well plates, and cells were overlaid with 1.5 ml DMEM (Invitrogen) supplemented with 0.8% methyl cellulose, 2% FBS, and 50 μg/ml gentamicin sulfate. Cells were incubated for 5 days at 37°C, fixed with 4% paraformaldehyde, and stained with crystal violet for plaque visualization.

### ZIKV RNA detection by *in situ* hybridization

Placental tissues from day 6 post infection were fixed in 10% neutral buffered formalin for 24 hours and placed back into PBS until paraffin embedding. *In situ* hybridization using RNAscope^®^ was performed on 5 μm paraffin-embedded sections. Deparaffinization and target retrieval were performed using RNAscope^®^ Universal Pretreatment Reagents (ACD #322380) following the manufacturer’s protocol, and fluorescence *in situ* hybridization was subsequently performed according to the manufacturer’s protocol (ACD# 323110) with RNAscope^®^ Probe V-ZIKVsph2015 (ACD #467871) as previously described [56]. Following *in situ* hybridization, slides were mounted with Vectashield hard-set mounting medium with DAPI (Vector Laboratories) and analyzed using an AxioImager Z2 microscope (Zeiss) and Zen 2012 software (Zeiss).

### Drug sensitivity assays

Vero cells (10,000 cells/well in a 96-well plate) were pretreated with media containing a carrier control or drug (ribavirin, mycophenolic acid, brequinar (Sigma) or atovaquone (ABCAM)) for 2 hours at 37°C. Following pre-incubation, cells were incubated with ZIKV or CHIKV-ZsGreen at a multiplicity of infection (MOI) of 0.1 in the presence of each drug or carrier control for 1 hour at 37°C. Cells were then washed extensively and media containing drug or carrier was added to each well. After incubation at 37°C for 48 h, cells were fixed with 4% paraformaldehyde and subject to immunostaining or visualized directly in the case of CHIKV-ZsGreen. In brief, following fixation the cells were washed with Perm-Wash buffer (BD Bioscience), permeabilized with 0.25% Triton-X 100 in phosphate buffered saline (PBS) (Gibco), and blocked with 0.2% bovine serum albumin (BSA) and 0.05% Saponin in PBS for 1 hour at room temperature (RT). Cells were then incubated with a monoclonal mouse antibody to the Flavivirus envelope protein (4G2) (Millipore) for 1 hr at RT. Following primary antibody incubation, cells were washed with Perm-Wash buffer and incubated with a secondary anti-mouse IgG antibody conjugated to Alexa488 for 1 hour at RT. DAPI staining protocol. Cells were then washed and infected cells were quantified on a CellInsight CX7 High-content microscope and screening platform using uninfected cells as a negative control and a cut-off for three standard deviations from negative to be scored as an infected cell.

To address the effect of atovaquone on infectious virion production, mammalian and insect cells were seeded as described above and infected with ZIKV at an MOI of 0.1 in the presence of increasing concentrations of atovaquone for 1 hr at 37°C. Cells were washed with PBS and incubated in media containing atovaquone or DMSO as a control for 36 hrs at 37°C. Virus containing supernatants were collected and viral titers were quantified by plaque assay.

### Cell viability assays

Cell viability was measured using the CellTiter 96 non-radioactive cell proliferation assay (Promega), according to the manufacturer’s protocol. Vero, 293T, and JEG3 cells (10,000 cells/well in a 96-well plate) were treated with increasing concentrations of atovaquone and incubated for 36 hours at 37°C. Following the incubation, 15 μL of dye solution was added to each well and incubated for 4 hours at 37°C with 5% CO2. The reaction was stopped by the addition of 100 μL of Solubilization/Stop Solution and the absorbance was measured at 570 nm in an EnVision microplate reader. The 50% cytotoxic concentration (CC_50_) was calculated by a non-linear regression analysis of the dose-response curves. Cell viability was also assayed using Sytox Green (Invitrogen) following the manufacteurer’s instructions. Sytox green positive cells were quantified on the CellInsight CX7 high-content microscope as described above.

### RNA extractions and RT-qPCR

Intracellular viral RNA was extracted with TRIzol reagent (Invitrogen) following the manufacturer’s instructions and used directly for cDNA synthesis with the Maxima H minus-strand kit (Thermo). Relative viral RNA levels were quantified using Power SYBR Green (Applied Biosystems) using the following primers [81]: GAPDH ((5’- GAAGGTCGGAGTCAACGGATTT −3’ and 5’- GAATTTGCCATGGGTGGAAT −3’) and ZIKV (5’- AGATGACTGCGTTGTGAAGC-3’ and 5’- GAGCAGAACGGGACTTCTTC-3’).

### Virus entry and time-of-addition assays

To assay for the role of atovaquone in viral entry, 293T cells were infected with ZIKV at an MOI of 0.1 in the presence of 4.5 μM atovaquone for 1 hour at 37°C. Cells were washed extensively, complete media with or without 4.5 μM atovaquone was added back to the cells, and virus containing supernatants were collected 36 hours post infection. Viral titers were quantified by plaque assay. As a control, media containing atovaquone was added after infection.

For time-of-addition studies, 293T cells, pretreated with 4.5 μM atovaquone for 1 hour or left untreated, were infected with ZIKV at an MOI of 0.1 in the presence or absence of atovaquone for 1h at 37°C. Following incubation, cells were washed to remove unabsorbed virus and media with or without atovaquone was added. At different time points post-infection, media was removed and media containing 4.5 μM atovaquone was added to the infected cells. Culture medium was collected at 36 hours post infection and viral titers were quantified by plaque assay. Media containing DMSO was used as a control for all time points.

### Rescue assay

Vero, 293T, JEG3, and C6/36 cells were seeded in a 96-well plate as described above. Cells were infected with ZIKV diluted to an MOI of 0.1 in DMEM containing atovaquone for 1h at 37°C. Following incubation, cells were washed three times with PBS to remove unabsorbed virus. After wash, cells were incubated with media containing atovaquone with either DMSO, or 100 μM uridine, cytidine, adenosine, or guanosine for 36hrs. Culture medium was collected at 36 hours post infection and viral titers quantified by plaque assay.

### Data analysis and statistics

GraphPad Prism 7.0 software was used for all analyses. The equations to fit the best curve were generated based on R2 values ≥ 0.9 [inhibitory concentration vs normalized response]. Two-way ANOVA and Students t-tests were also used, with P values <0.05 considered statistically significant.

## ACKNOWLEDGMENTS

We thank all members of the Stapleford lab for support and helpful discussions during the course of the study. We thank Aaron Briley and Meike Dittmann for technical assistance with the CX7 CellInsight High content microscopy and helpful discussion. Finally, we thank the labs of Drs. Carolyn Coyne, Matthew Evans, and Alexander Pletnev for essential cells and reagents.

## Figure Legends

**Figure 2: Atovaquone inhibits ZIKV infectious virion production in mammalian cells.**

(**A**) Vero, (**B**) 293T, and (**C**) JEG3 cells were infected with ZIKV MR766 at an MOI of 0.1 in the presence of atovaquone (open symbols) or DMSO control (closed symbols). Virus containing supernatants were collected 36 hours post infection and infectious virus was quantified by plaque assay on Vero cells. The mean and SEM are shown, n=9, Students *t*-test, * p<0.05.

**Figure 3: Atovaquone inhibits chikungunya virus replication.**

(**A**). Vero cells were pretreated with DMSO or atovaquone (ATQ) for two hours and subsequently infected with CHIKV (IOL) expressing ZsGreen at a MOI of 0.1. After infection, virus was removed, cells washed and media containing atovaquone was added for 24 hours. Cells were then fixed and ZsGreen positive cells were quantified by a CellInsight CX7 high-content microscope. Data are represented as percent ZsGreen positive compared to the DMSO control. (**B**). 293T and Vero cells were infected with CHIKV ZsGreen at a MOI of 0.1 in the presence of 4.5 μM atovaquone. Unabsorbed virus was washed off and media was added containing DMSO or 4.5 μM atovaquone. Infectious virus was quantified 24 hours post infection by plaque assay. The mean and SEM are shown, n=3, Students *t*-test. *** p<0.005.

**Figure 4: Atovaquone acts early during ZIKV infection and inhibits viral RNA synthesis.**

(**A**). 293T cells were infected with ZIKV MR766 at an MOI of 0.5 in the presence of 4.5 μM atovaquone. After absorption, cells were washed extensively, and media was added without (at entry) or with (post entry) 4.5 μM atovaquone. Infection virus was quantified by plaque assay 36 hours post infection. The mean and SEM are shown, n=4, Students t-test, * p<0.05. (**B**). 293T cells were treated with 4.5 μM atovaquone one hour prior, during, or post infection with ZIKV MR766 at a MOI of 1. After infection, virus was removed and media containing 4.5 μM atovaquone was added for 36 hours. Infectious virus was quantified by plaque assay 36 hours post infection. The mean and SEM are shown, n=3, Students t-test, * p<0.05, ** p<0.01 (**C**). Vero cells were infected with ZIKV MR766 at a MOI of 1 in the presence of 4.5 μM atovaquone. After infection, virus was removed, cells washed extensively, and media was added containing 4.5 μM atovaquone. At time 0 (after infection), 24, and 48 hours post infection, media was removed, cells washed, and intracellular RNA extracted with Trizol. Viral RNA was quantified by SYBR Green compared to GAPDH. The mean and SEM are shown, n=3, Students t-test, **** p<0.0001.

**Figure 5: Exogenous uridine rescues ZIKV virion production and RNA synthesis.**

(**A**) 293T, (**B**) JEG3, and (**C**) Vero cells were infected with ZIKV MR766 at a MOI of 0.1 in the presence of DMSO, atovaquone (293T and Vero = 4.5 μM atovaquone and JEG3 = 18 μM atovaquone), or atovaquone with 100 μM of uridine (U), cytidine (C), adenosine (A), or guanosine (G). After infection virus was removed, cells washed, and media was added containing DMSO, atovaquone, or atovaquone with 100 μM nucleoside. Infectious virus was quantified 36 hours post infection by plaque assay. The mean and SEM are shown, n=3, two-way ANOVA, * p<0.05, *** p<0.005, **** p<0.0001. (**D**). Vero cells were infected with ZIKV MR766 at a MOI of 1 in the presence of DMSO, 4.5 μM atovaquone, or 4.5 μM atovaquone with 100 μM uridine. Intracellular viral RNA was extracted with Trizol at 0, 24, and 48 hours post infection and viral RNA relative to GAPDH was quantified by Sybr Green. The mean and SEM are shown, n=3, two-way ANOVA, **** p<0.0001.

**Figure 6: Atovaquone inhibits ZIKV infection and spread in an *ex vivo* human placental tissue model.**

Human placental chorionic villus explants were infected with 10^5^ PFU ZIKA (MR766) in the presence of increasing concentrations of atovaquone or a DMSO carrier control. Supernatants were collected at days 2, 4, and 6 post infection and viral titers were quantified by plaque assay. (A) Total virus accumulation over the course of infection. The mean and SEM are shown, n=3, two-way ANOVA, *** p<0.005. (B) Fluorescence *in situ* hybridization of ZIKV RNA (red) infected human tissues counterstained with DAPI at 6 days post infection.

